# The modelling of community assembly during seagrass restoration

**DOI:** 10.64898/2026.02.24.707629

**Authors:** Jane Allwright, Jim Bull, Mike Fowler

## Abstract

Successful seagrass restoration will provide habitat for a variety of species. Here, ecological community assembly in a newly planted seagrass meadow has been modelled mathematically using a combination of numerical integration and a permanence-based method, and using real data to parametrise the models. We have studied the transient dynamics of the system: how the ecological communities assemble and change over a 100-year period. Using a trophic structure and a range of species pool sizes, we investigated how much variability there was in community size for a given sized species pool, whether it is possible to use early monitoring to predict the final community size, and to what extent monitoring gives an indication of final vs transient species. For the majority of cases modelled, the community either reached or was headed towards an endpoint community which was uniquely determined by the species pool. However, for 1.4% of cases, no unique endpoint community could be calculated. The simulated communities began to assemble within the first ten years, but 13% had still not reached their endpoint community even after 100 years. In 62% of our models, no consumer species colonised in the first two years, suggesting that monitoring should certainly be continued beyond a two-year period. We counted how many of the species that were present at any observation point in the 100 years would also be present in the endpoint community, and found that this proportion generally decreased with increasing species pool size, to an average of 86% when the species pool had 49–56 consumer species. By monitoring the community over the first ten years, it is not possible to deduce what the final community will be; however a very small number of fauna species present over the first ten years might be used to predict very small endpoint communities.

## 1 Introduction

We have used mathematical modelling techniques to study ecological community assembly in restored seagrass meadows. Seagrass restoration projects are being carried out globally [1, 2, 3] and around the British Isles [4, 5], with seagrass known to have many benefits such as carbon storage, coastal protection, and providing habitat for a variety of species [6]. As a seagrass meadow becomes established and develops over time, both the number of fauna species living in it (species richness) and the abundance or biomass of each one will vary as the different species interact with one another. This raises the question of how these communities change over time. We have therefore looked into the transient dynamics of the system, as well as the asymptotic or theoretical long-term states.

We have applied numerical integration of Lotka-Volterra systems, together with the permanence-based modelling approach of [7, 8, 9], to the specific ecological example of seagrass restoration. Our study has focused on the Zostera marina species of seagrass, and on restoration around the coasts of the British Isles. However our modelling approach was designed to be sufficiently general that, by changing parameter values and trophic structure, it can readily be adapted to other habitat restoration scenarios. The aims of this project were to increase understanding of community assembly, improve predictions, and help inform monitoring programmes.

The need for long-term monitoring is becoming well recognised. In a review of seagrass restoration in Australia and New Zealand [10], Tan et al. report that although one restoration site in that area had been monitored for over 20 years, more than a third of restoration projects in Australia and New Zealand did not monitor beyond a year. They therefore emphasise the need for long-term monitoring. They also recommend a programme of regular monitoring, as opposed to simply monitoring at the start and end of the restoration trial. The handbook for seagrass restoration in Scotland [11] also highlights the need for a well-designed monitoring programme, and one which takes account of the aims of the project. It says that “seagrass growth from seed takes as little as 12 months, but development into mature bed takes many years and the knock-on ecosystem effects may not be measurable for a decade or so”. Therefore it recommends that the monitoring of seagrass restoration projects in Scotland should continue for a minimum of ten years, and it describes the point six years after planting as a “key time point for assessing restoration success” [11]. Orth et al. [12] agree that a “sustained commitment to long-term monitoring and research” is “essential”, and that reasonable timescales for the reversal of seagrass ecosystem degradation are in the order of years to decades. In their study into Zostera marina restoration in the mid-western Atlantic (Virginia, USA) they report that epifaunal invertebrate communities containing crustaceans, decapods and gastropods in the restored areas were “indistinguishable” from nearby natural communities by 3 years after restoration. They comment that “faunal response to the restoration effort was initially marked, with values rapidly matching those from other areas and saturating in less than a decade” [12].

When seagrass restoration is successful, the planted seagrass grows to form meadows which provide habitat for a variety of species. Several field studies have been carried out on ecological communities in Zostera marina habitats [13, 14, 15, 16]; these provide useful information about the typical sizes and structures of the communities that live in seagrass meadows. The biodiversity of fauna in terms of species richness and species abundances have been measured and, in some cases, monitored over time. In [13] it was recorded how the number of taxa, number of individuals and diversity all increased with seagrass shoot density and with time. The total number of taxa found in four separate studies [13, 14, 15, 16] and the available information regarding average community size are summarised here in Table 1. These suggest that realistic models should have species pools of between about 23 and 124 animal species, and should typically produce communities consisting of 5 to 10 species — although the observed community size was not reported in some instances. These studies also gave information about which taxa were dominant, and [14, 15, 16] included lists of the species that were identified.

**Table 1:** Reported species richness in field studies of Zostera marina.

| Reference | Location | Total number in study | Average community size |
| --- | --- | --- | --- |
| [13] | North Carolina, USA | 89 taxa | — |
| [14] | Northern Baltic sea | 37 species | 5.9–8.8 species |
| [15] | Ireland | 124 species, representing<br>81 families from 6 phyla | — |
| [16] | Atlantic coast of Canada | 23 species or genera | 7–10 species |

Our model used a multi-trophic food web structure. We used the species lists from [15] as a starting point and then grouped them together according to prey type until we had a sufficiently simple model formed of four trophic levels. The first (basal) trophic level was the primary producer and can be thought of as including the seagrass itself and also the epiphytes growing on it, other algae, detritus and phytoplankton. Trophic level 2 contained grazers, herbivores, detritivores, suspension feeders and zooplankton. For a seagrass community in the British Isles this would include, for example, shell fish, small crustaceans and other small invertebrates, sea urchins, zooplankton, gastropods, and herbivorous fish or marine worms. Trophic level 3 — the mesopredators — would include starfish, carnivorous marine worms, anemones, and some predatory fish and crustaceans. The top level of the model, level 4, would contain predators such as crabs and sea spiders which may take their prey from both the mesopredator level and the herbivore level (trophic levels 3 and 2).

Using this food web structure, we considered a total of 1600 species pools across a range of sizes. Through our modelling, we investigated how communities assemble and change over time. For each case, we studied how the community varies both in species richness and in composition, whether it reaches or tends towards an endpoint community, the timescales involved, and whether the final community is determined solely by the species pool or whether other factors (such as the order of species arrival) may be important.

We had three key research questions. Beginning with the set of all seagrass meadows and asking how much consistency or variability there was between them, our first question was: how variable is community size for a given sized species pool? Secondly we asked: for each individual seagrass meadow, to what extent is it possible to predict final community size based on an initial period of monitoring? In other words, is it possible to use early monitoring to predict the number of species in the final community, and if so, after how many years does it become possible? Our third question was related to the individual species identities in the community: to what extent does monitoring give an indication of the final species vs the transient species?

## 2 Methods

### 2.14 Lotka-Volterra model

As in [23], our model was based on a Lotka-Volterra ODE system,

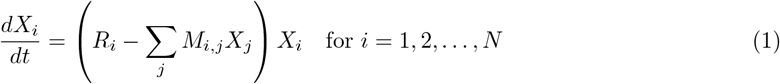

where *X*_*i*_(*t*) were the biomass densities of each species (*i* = 1, 2, …, *N*), *R*_*i*_ were the species-specific intrinsic growth/mortality rates, and the matrix *M* contained the intra- and inter-specific interaction coefficients. The inter-specific interactions (the off-diagonal elements of *M*) were of predator-prey type, with each species assigned to a certain trophic level in the food web and consuming and/or being consumed by other species. Thus *M*_*i,j*_ < 0 corresponded to species *i* consuming species *j*, and *M*_*i,j*_ > 0 for species *i* being consumed by species *j*. The model also included a small amount of intra-specific competition, with diagonal entries *M*_*i,i*_ > 0, but we did not include any direct inter-specific competition.

Our model food webs contained four trophic levels. Level 1 consisted of a single species (or functional group), *X*_1_, representing the primary producer. This was the only species with a positive *R*_*i*_ value. All of the other species were consumers with *R*_*i*_ < 0. Each of the animal species *X*_2_, … *X*_*N*_, was assigned to a particular trophic level. The herbivores (trophic level 2) consumed the plant species and were themselves consumed by species in higher trophic levels. Mesopredators in trophic level 3 consumed prey from level 2, while trophic level 4 consisted of ‘apex predators’ which could consume prey from both level 3 and level 2. We modelled species pools of a range of sizes (*N*) from 8 to 57 species. The number of apex predators varied from 1 to 8 and we always assumed a pyramidal food web of consumers, with twice as many species in level 3 as in level 4, and twice as many species in level 2 as in level 3 (see Table 2). We used random sampling of *R*_*i*_ and *M*_*i,j*_ values to construct 200 different species pools of each size (see Section 2.1.1 for details on how the *R*_*i*_ and *M*_*i,j*_ were chosen).

**Table 2:** Number of species modelled in each trophic level.

| Level 1 (plant) | Level 2 (herbivores) | Level 3 (mesopredators) | Level 4 (apex predators) | Total $N$ |
| --- | --- | --- | --- | --- |
| 1 | 4 | 2 | 1 | 8 |
| 1 | 8 | 4 | 2 | 15 |
| 1 | 12 | 6 | 3 | 22 |
| 1 | 16 | 8 | 4 | 29 |
| 1 | 20 | 10 | 5 | 36 |
| 1 | 24 | 12 | 6 | 43 |
| 1 | 28 | 14 | 7 | 50 |
| 1 | 32 | 16 | 8 | 57 |

#### 2.1.1 Interaction coefficients

There are several sources of data available on the biomass density and production rates of seagrass; see for instance [17, 18, 19, 20]. Measured values for seagrass above-ground biomass are typically a few hundred grams (dry weight) per square metre in a seagrass meadow, and the rate of leaf biomass production is a few grams per square metre per day. In [18] Duarte and Chiscano reported that, averaged over several seagrass species, the maximum above-ground biomass was 223.9 ± 17.5*g*/*m*^2^, and the above-ground biomass production was 3.84 ± 0.34*g*/*m*^2^/*day*. In Table 3 we have summarised the data from [17, 18, 19, 20] for the particular species Zostera marina. We used this to choose the coefficients *R*_1_ and *M*_1,1_ in a suitable range, given that the units used in our models were *g*/*m*^2^ and days. For each new each species pool that we generated, we took a maximum plant production rate *r* from a uniform distribution on the range [2.5, 7.9]*g*/*m*^2^/*day* and a carrying capacity *K* = *K*_0_ × 1.3 where *K*_0_ was chosen from a uniform distribution on the range [280.9, 315.9]*g*/*m*^2^. The factor of 1.3 represented the fact that the variable *X*_1_ in our model should incorporate not only the seagrass but also the epiphytes — which are reported as contributing an extra biomass of around 30% of that of the seagrass itself [21, 22]. The value of *K* and the rate *r* were included in our model by choosing

**Table 3:** Reported biomass density and production rates of Zostera marina. (The values reported in [20] were the mean vegetative shoot biomass and reproductive shoot biomass; we give the sum of these values.)

| Reference | Location | Reported above-ground biomass | Reported above-ground (leaf) biomass production | Average above-ground (leaf) biomass production per day |
| --- | --- | --- | --- | --- |
| [17] | Denmark | $38 - 219g/m^2$ | $654 - 995g/m^2$ over a year | $1.79 - 2.72g/m^2/day$ |
| [19] | Denmark | max. $443g/m^2$ | $856g/m^2$ over 190 days, and max. $7.9g/m^2/day$ | $4.5g/m^2/day$, and max. $7.9g/m^2/day$ |
| [18] | a range of locations | $298.4g/m^2$ | $5.2g/m^2/day$ | $5.2g/m^2/day$ |
| [20] | China | $366.5g/m^2$ (2009), $321.0g/m^2$ (2012) | — | — |

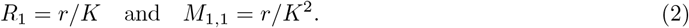

This ensured that the carrying capacity of the plant species was *R*_1_/*M*_1,1_ = *K* and that, at its carrying capacity, the production rate of the plant would be *R*_1_ × *K* = *r*.

Each consumer species had *R*_*i*_ chosen from a uniform distribution on the range [−0.1, −0.01] representing mortality rates of one every 10 to 100 days. As in [23], the values were sorted so that the death rates decreased (*R*_*i*_ got closer to zero) as we moved up the food web from herbivores to the higher trophic species (i.e. species in higher trophic levels tend to live longer).

Each consumer species was assigned a small, random intra-specific competition value, with *M*_*i,i*_ taken from a uniform distribution on the range [0.005, 0.05]. There was no direct inter-specific competition.

Each consumer species *X*_*i*_ had a conversion efficiency *c*_*i*_ which was chosen from a uniform distribution on the range [0.08, 0.12]. This meant that if species *i* consumed species *j* then

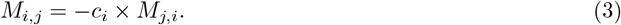

For each herbivore *X*_*i*_, the rate at which it consumed the plant was chosen large enough that it would be able to survive in the 2-species system consisting only of the plant and the herbivore (*X*_1_ and *X*_*i*_). This 2-species system would have a positive equilibrium provided that

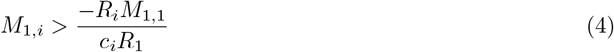

or equivalently, using equation (2), 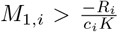. In our model, for each herbivore *X*_*i*_ the coefficient *M*_1,*i*_ was chosen from a uniform distribution on 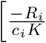, max 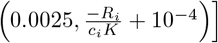.

For each predator *X*_*i*_ (i.e. in trophic level 3 or 4), the number of prey species it would consume from the trophic level beneath it was chosen between 1 and the total number *n* in that level. For *m* = 1, …, *n*, the number of prey species was chosen as *m* with probability 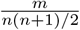. That is, predators were more likely to be generalist than specialist. The *m* prey species were then chosen at random from the relevant trophic level. For each such prey species *j*, the coefficient *M*_*j,i*_ was chosen from a uniform distribution on [0.2, 0.4].

We allowed apex predators (in trophic level 4) to also take prey from trophic level 2. For each species *X*_*i*_ in trophic level 4, the number of prey species it would consume from level 2 was chosen between 1 and half the total number *n*_2_ in level 2. Each number from 1 to 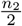 was equally likely, and that number of prey species was then chosen at random from level 2. Again, for each such prey species *j, M*_*j,i*_ was chosen from a uniform distribution on [0.2, 0.4].

#### 2.1.2 Simulation, initial conditions, and threshold

For each species pool, we used the numerical ODE solver ‘ode45’ in MATLAB R2022b to solve the population dynamics. We simulated each model for 36525 days, i.e. 100 years. The species densities were recorded at the end of each simulated year (365.25 days) to allow us to evaluate how the community changed from one year to the next.

As we were interested in community assembly on a newly planted seagrass restoration site, we used ‘small’ initial conditions of 5*g*/*m*^2^ for the plant species (representing the first seedlings) and 5 × 10^−5^*g*/*m*^2^ of each other species.

Since all species in the species pool start with a positive density *X*_*i*_ at time *t* = 0, it follows from the Lotka-Volterra equations (1) that they remain strictly positive for all *t* > 0. However some of the densities *X*_*i*_ may in fact be very close to zero. We defined ‘the community’ at the end of each simulated year to be the set of species with densities above a fixed threshold value of 10^−4^*g*/*m*^2^. These are also referred to as the species ‘present’ at the end of each simulated year. Note that the initial densities of consumer species were deliberately below this threshold, since one would expect all consumer species to be absent until after the seagrass had begun to grow.

One question we were interested in was how long it would take for the number of consumer species present in the community to go above, and then stay above, zero (i.e. for the seagrass to be colonised). Once there was at least one consumer present, we also monitored the diversity measures *N*_0_, *N*_1_ and *N*_2_ (using the notation of Hill [24]) over time. *N*_0_ is the total number of consumer species present in the community (i.e. those above the threshold), *N*_1_ is equal to exp(*H*) where *H* is the Shannon diversity of the consumer species present in the community, and *N*_2_ is the reciprocal of Simpson’s index for these consumer species.

### 2.2 Uninvadable and permanent communities

In addition to investigating how each community developed over time, we were interested in knowing whether it ever reached, or was heading towards, a ‘final’ (endpoint) community. For such cases, we were interested in the timescales over which these endpoint communities were reached, and what they were like in terms of number of species from each trophic level and the total number of species. To determine the endpoint communities, we used the concepts of uninvadability and permanence [7, 8, 9].

#### 2.2.1 Permanence, uninvadability, and average Lyapunov functions

Consider a subset *s* of *n* species, taken from a species pool of total size *N*. The subset *s* has a positive equilibrium if there exists a fixed point *X*^(*s*)^ of the equations (1) with a positive density for each species in *s* and zero density for all other species from the pool:

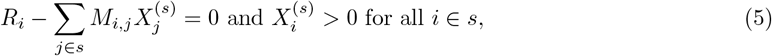

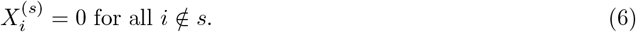

If a subset *s* has a positive equilibrium, we say that it is ‘uninvadable’ if none of the other *N* − *n* species from the species pool is able to invade *X*^(*s*)^. Mathematically, this corresponds to the condition

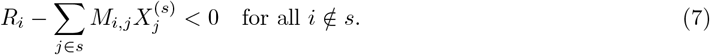

Conversely, if any species not in *s* has a positive growth rate near to *X*^(*s*)^, that is

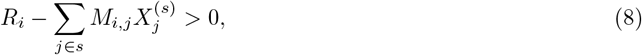

then this species can invade *s*.

A subset *s* of species is said to be ‘permanent’ if the species will coexist with each other, without any either growing indefinitely or tending towards extinction. More precisely, if the species in question all start with strictly positive densities, then they will remain both bounded above and bounded below some fixed distance away from zero. The permanence condition makes no assumption about the asymptotic behaviour (a permanent subset might have, for example, an asymptotically stable equilibrium point, a limit cycle, or a chaotic orbit); it simply means that the species will coexist together over the long-term [25, 7, 8, 9, 26]. In our models, the species’ densities always have finite upper bounds. Therefore, to check whether a subset *s* satisfies the permanence condition, we want to know whether the boundary of its phase space (where one or more of the species in *s* has zero density) repels trajectories, thereby keeping them in the interior. In [7, 8, 9] it is explained that, due to the Lotka-Volterra form of the equations (1), and results of Hofbauer and Sigmund [27] and Jansen [28], the question of permanence can be linked with the existence of an ‘average Lyapunov function’. Moreover, a linear optimisation problem can be used to establish whether there exist a choice of positive exponents *h*_*i*_ > 0 such that the function

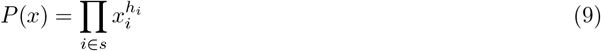

is an average Lyapunov function (see Appendix A). If there is such a choice of the *h*_*i*_ then the subset *s* does satisfy the permanence condition. While this method provides a sufficient condition for permanence, it is not a necessary condition [7, 8, 9]. In other words, it is possible for a subset *s* to be permanent and yet not possess an average Lyapunov function of the form (9). This means that this test could fail to recognise certain permanent subsets.

#### 2.2.2 Identifying the endpoint community

To identify the endpoint community (or communities) of each species pool, we searched for permanent and uninvadable subsets. If the species pool was small enough (*N* ≤ 25 species), we tested all 2^*N*^ possible subsets for positive equilibria, uninvadability (7), and the permanence condition (by solving the linear programming problem using the ‘linprog’ function in MATLAB R2022b). If there was exactly one subset that had a positive equilibrium, was uninvadable, and satisfied the sufficient condition for permanence, then this was considered the endpoint (‘final’) community.

As computational time increased prohibitively for *N* > 25 with this approach, we instead used an iterative method similar to [9, 29]. We began with the empty subset and, since the plant species *X*_1_ was the only species with positive intrinsic growth rate, we ended the first iteration with the subset consisting of just *X*_1_. Each other iteration began with the community (subset of species) determined by the preceding iteration. On each iteration, starting from a community *s* of *n* species, we added another species chosen at random from the set of those satisfying (8) (i.e. the set of species from the species pool that could ‘invade’ successfully). Then we took this temporary set of size *n* + 1 and tested all 2^*n*+1^ possible subsets for positive equilibria, permanence (using the sufficient condition as discussed above), and uninvadability. If there was exactly one subset that had a positive equilibrium, was uninvadable, and satisfied the sufficient condition for permanence, then this became the community from which to start the next iteration. If there was more than one such subset then numerical integration was used, starting from the positive equilibrium of the original *n* species plus a small initial value (10^−4^) of the invading species. Those species that were above the threshold density after simulating for 200 years then became the next community from which to start the next iteration (after checking that set did indeed have a positive equilibrium). If, on the other hand, the temporary set of size *n* + 1 had no subsets that had positive equilibria, were uninvadable, and satisfied the sufficient condition for permanence, then we searched for subsets that had positive equilibria, were permanent and contained the new (invading) species. From the collection of such subsets, we found those subsets having the largest size, and one of these was then chosen at random; this became the community from which to start the next iteration.

If this iterative process reached a state that was uninvadable by all remaining species, this was considered a ‘terminal’ community and the iterations stopped. Otherwise, iterations continued up to a maximum number of 200 iterations. We repeated the iterative method 20 times per species pool, thereby finding up to 20 possible terminal communities. If all 20 runs of the iterative method reached the same terminal community, then this was considered the endpoint community for that species pool.

There were a small number of cases where no endpoint community could be found by the above process. This included species pools where there was more than one permanent and uninvadable community. It could also include species pools that had no subsets that were permanent and uninvadable, species pools where all such subsets failed the test for permanence (which is only a sufficient condition but not necessary), or species pools where certain permanent and uninvadable subsets either took more than 200 iterations to reach or could not be reached at all by the method of sequential invasion.

For those cases that did have a unique endpoint community, we measured the size of this community (the number of consumer species) and its composition (how it was split across the trophic levels).

#### 2.2.3 Comparison of simulated community and the endpoint community

For the cases that had a unique endpoint community (as identified using the method in Section 2.2.2), we wanted to know whether the simulation actually approached this state over time, and if so then over what timescale it was reached.

One method we used to study this was to simply consider species richness: when does the size *N*_0_ of the community approach that of the endpoint community? And, to what extent can the species richness over the first ten years be used to predict high or low species richness of the endpoint community?

In order to look into whether the species present at different points of the simulation matched those in the endpoint community or were transient species, we looked at the number of species ‘missing’ (that is, species that were absent at a given point in time but were present in the endpoint community) and the number of species that were ‘extra’ (that is, species that were present at a given point in time but were not present in the endpoint community). The community from the simulation thus approaches the endpoint (‘final’) state if and only if the number of ‘missing’ species and the number of ‘extra’ species both go to zero.

We also asked, out of all the species observed at any year throughout the simulation, what proportion of them were present in the endpoint community.

## 3 Results

### 3.1 Modelling results

In the initial conditions, all consumer species had densities below the threshold value and were therefore absent from the community. After a certain time the number of consumer species present went above zero (i.e. the seagrass was colonised). From then on, the number of consumer species present varied over time; see Figures 1 and B1. On average, the number *N*_0_ of consumer species present increased over time and generally levelled out at a constant value (see Figures 1 and B1). But for individual models, *N*_0_ did not monotonically increase; it sometimes went down as well as up.

**Figure 1.**
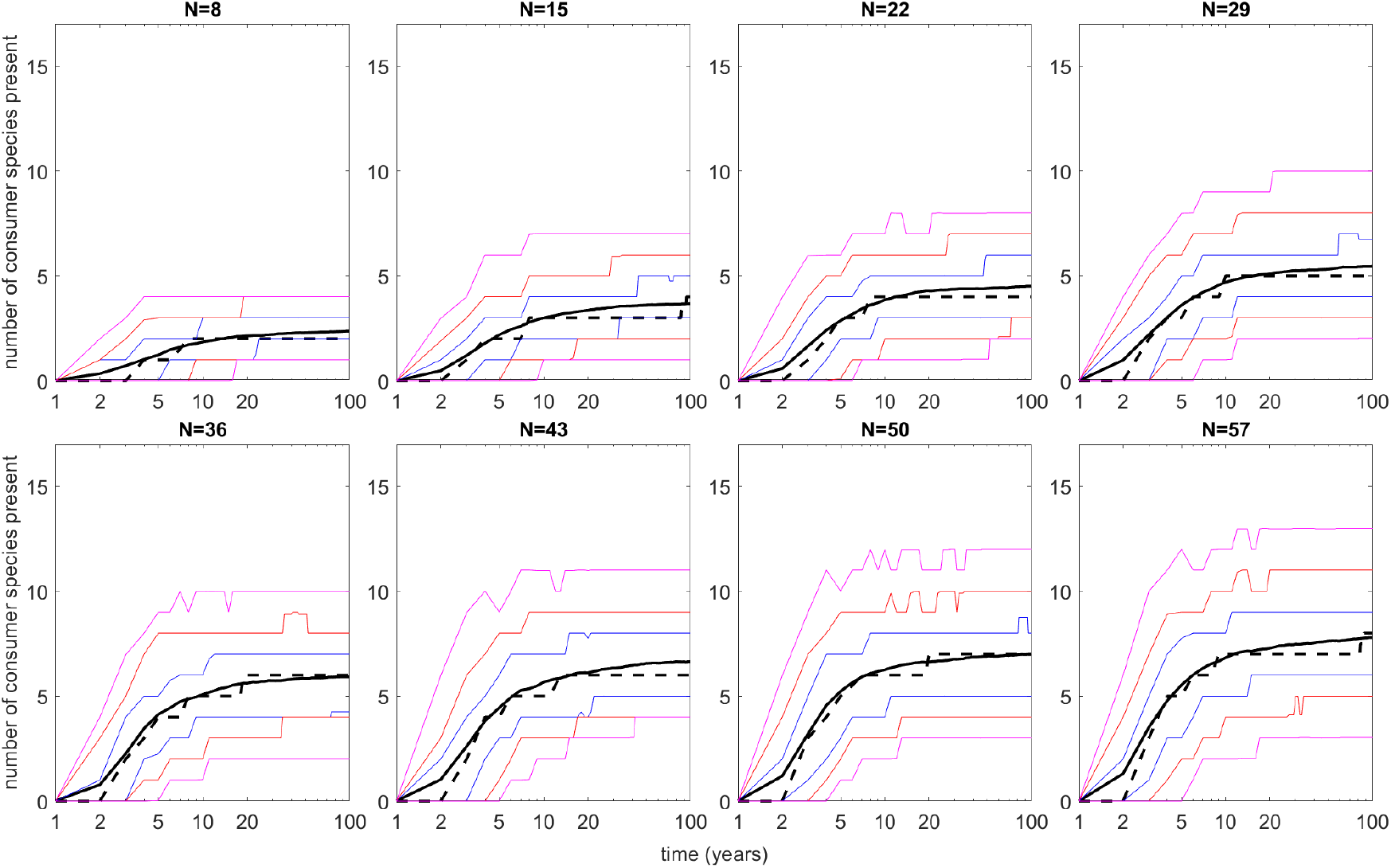
The number of consumer species present in the community, over time. The black solid line shows the mean and the black dashed line the median. The blue lines indicate the 25th and 75th percentiles, the red lines indicate the 10th and 90th percentiles, and the pink lines indicate the 2.5th and 97.5th percentiles. Therefore the middle 50% of cases lie between the blue lines, the middle 80% between the red lines and the middle 95% between the pink lines. Time is plotted on a logarithmic scale.

We found that by two years, 62% had not yet been colonised, while 38% of the cases had at least one consumer species present and 22% of cases had two or more consumers present (see Table B1). From five years onwards, 91% of the cases had at least one consumer present, and 78% had at least two consumer species present. As one would expect, the proportions of cases with at least one or at least two species present were higher for the larger species pools and lower for the smaller species pools. By ten years, there were only 1.3% of cases remaining (21 out of 1600) that still had no consumer species present, and these were primarily from the very small (*N* ≤ 15) species pools.

We also plotted the diversity measures *N*_0_, *N*_1_ and *N*_2_ over time; see Figure B1. Note that, for the purpose of plotting, we set *N*_0_, *N*_1_ and *N*_2_ to be zero at the early stages of the simulation while no consumer species were yet present. After that they were defined according to Hill [24]. The diversity measures *N*_1_ and *N*_2_ behaved, qualitatively, very similarly to *N*_0_ (see Figure B1) and did not give much extra insight into the community assembly process.

For larger species pools (*N* ≥ 29 species), Figure B1 and Table 5 show that the typical community sizes *N*_0_ were in the range 5–10. This is the realistic range of values reported from field surveys (see Table 1).

The approach described in Section 2.2.2 was used to determine the endpoint community for each species pool. Out of the 1600 species pools generated in total (200 for each size *N*), there were 23 (1.4% of cases) for which the method did not find a unique endpoint community. Table 4 indicates how these were spread across the species pool sizes, and the different reasons for lacking a unique endpoint community. In total there were 14 species pools (0.88% of cases) with two permanent and uninvadable communities. In our study we did not find any cases with more than two permanent and uninvadable communities, although this is also possible in principle.

**Table 4:**
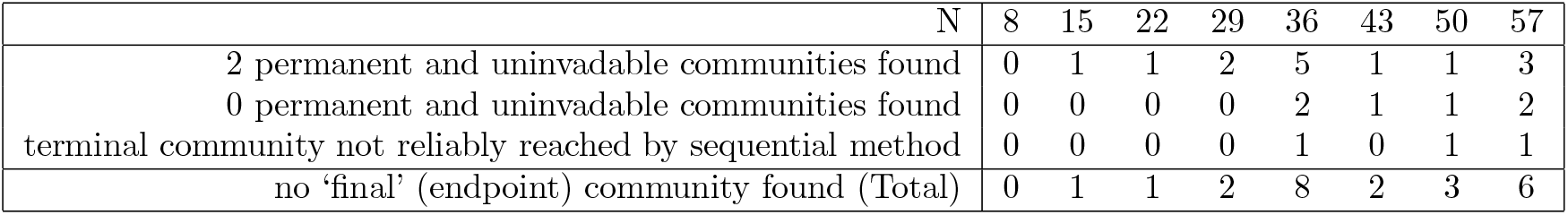
Number of cases (out of 200 species pools for each *N*) without a unique endpoint community.

**Table 5:**
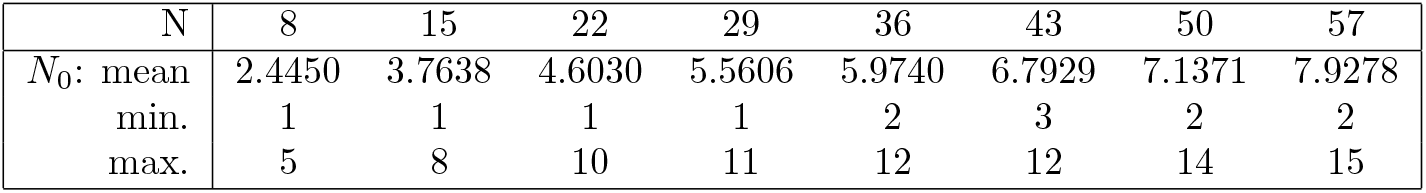
Number *N*_0_ of consumer species in the endpoint communities.

| N | 8 | 15 | 22 | 29 | 36 | 43 | 50 | 57 |
| --- | --- | --- | --- | --- | --- | --- | --- | --- |
| $N_0$: mean | 2.4450 | 3.7638 | 4.6030 | 5.5606 | 5.9740 | 6.7929 | 7.1371 | 7.9278 |
| min. | 1 | 1 | 1 | 1 | 2 | 3 | 2 | 2 |
| max. | 5 | 8 | 10 | 11 | 12 | 12 | 14 | 15 |

For each of the remaining 1577 species pools we did find a unique endpoint community. Moreover, at the equilibrium of the endpoint community, the positive densities were all above the threshold value of 10^−4^.

For the 1577 species pools that had a unique endpoint community, we measured the size of this community (the number of consumer species) and its composition (how it was split across the trophic levels); see Table 5 and Figure B2. The Spearman’s rank correlation coefficient between the total size *N* of the species pool and number of consumers in the endpoint community was 0.6836.

### 3.2 How much variability is there in community size for a given sized species pool?

In Figure 1, the middle 50% of cases lie between the blue lines, the middle 80% between the red lines and the middle 95% between the pink lines. This demonstrates the substantial variability in the community size throughout the assembly process.

The mean, minimum and maximum size of the endpoint community are given in Table 5 for each sized species pool. The standard deviation was between 28% and 40% of the mean endpoint community size. The distributions of endpoint community sizes, together with the numbers in each of trophic levels 2, 3 and 4, are shown in the histograms in Figure B2.

### 3.3 How useful is early monitoring to predict the endpoint community size?

The majority of our models (62%) did not yet have a consumer species present (above the threshold for observation) in the first two years (see Figure 1 and Table B1), suggesting that monitoring should certainly be continued beyond a two-year period.

By five years after the start of the simulation 91% of the models had at least one consumer present, and by ten years almost all cases had one or more consumers present. The few cases (1.3%) that still had no consumer species present by that point were primarily from the very small (*N* ≤ 15) species pools and are therefore less likely to be realistic for real seagrass communities. This suggests that the ecological community may start to form within the first five to ten years after the appearance of the seagrass seedlings.

For the 1577 cases that had a unique endpoint community, we monitored how *N*_0_ approached the number of consumer species in the endpoint community. See Figures 2 and B3 where the number of consumers present is plotted as a proportion of the number in the endpoint community.

**Figure 2.**
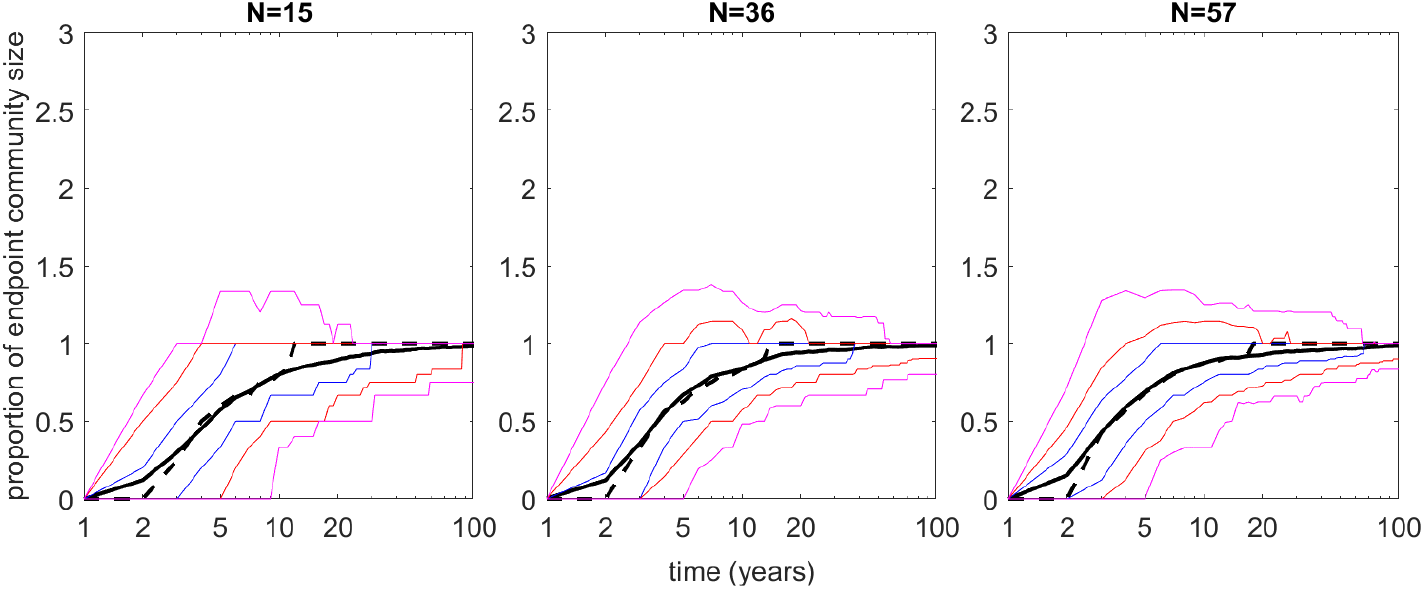
Number of consumer species present, as a proportion of the number in the endpoint community. The black solid line shows the mean and the black dashed line the median. The blue lines indicate the 25th and 75th percentiles, the red lines indicate the 10th and 90th percentiles, and the pink lines indicate the 2.5th and 97.5th percentiles. Therefore, the middle 50% of cases lie between blue lines, the middle 80% between the red lines, and the middle 95% between pink lines. Time is plotted on a logarithmic scale.

We also asked to what extent the number of consumer species present over the first 2–10 years could be used to predict small or large sizes of the endpoint community. For the 1577 cases that had a unique endpoint community, we plotted the number of consumers in the endpoint community firstly against the maximum consumer species richness from years 1–2, then against the maximum from years 1–5, and finally against the maximum from years 1–10 of the simulation. These are plotted in Figure 3, where the diameters of the dots increase with the number of cases having each combination. The Spearman’s rank correlation coefficient for the first case (years 1–2) was 0.3352, for the second case (years 1–5) Spearman’s rank correlation coefficient was 0.7339, and for the third case (years 1–10) it was 0.8520. By monitoring the community over the first ten years, it is not possible to deduce what the final community size will be. However there is a strong correlation between the maximum species richness from years 1–10 of the simulation and the size of the endpoint community. Figure 3 shows that the cases where no consumer species were present throughout years 1–10 (which was 21 cases out of 1577) also had a very low (≤ 4) species richness in their endpoint community, and those with a maximum of one consumer species over years 1–10 (which was 117 cases out of 1577) also had small endpoint communities, with ≤ 6 consumer species.

**Figure 3.**
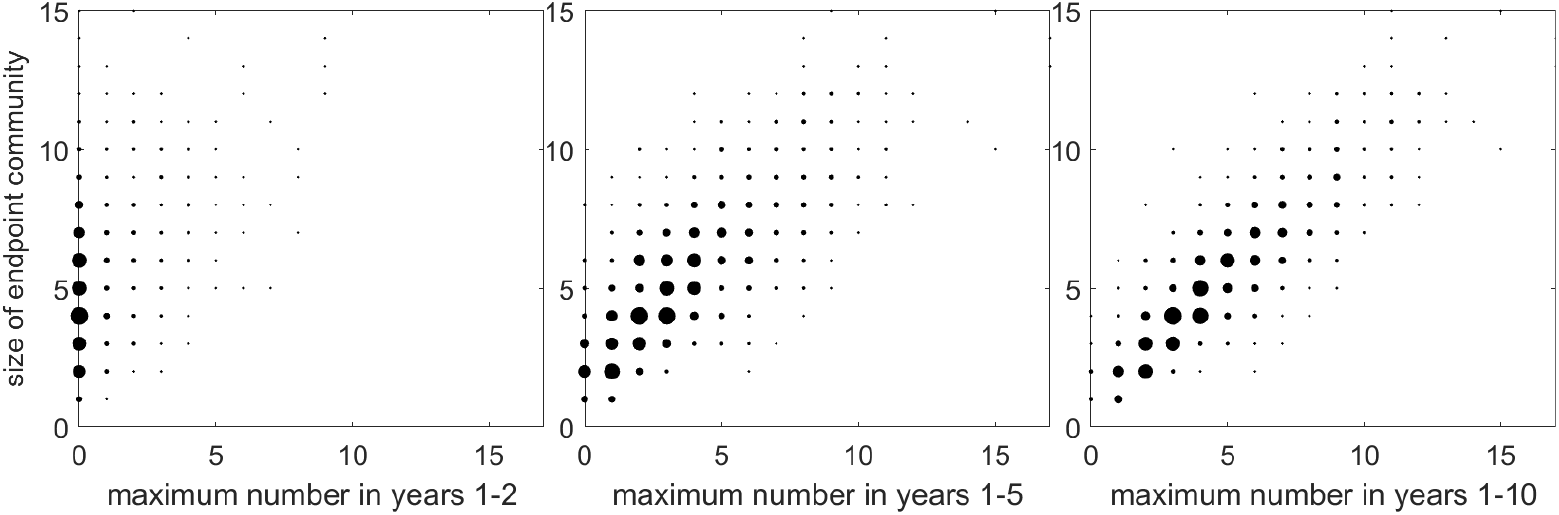
Number of consumer species in endpoint community vs the maximum over years 1–2, 1–5, and 1–10 of the simulation. Diameters of the dots are related to the number of cases having each combination.

### 3.4 How many of the observed species are ‘final’ vs transient?

For the 1577 cases that had a unique endpoint community, we counted the number of species ‘missing’ and the number of ‘extra’ species (relative to the endpoint community) over time. See Figures 4, B4, 6 and B6. Also, in Figures 5 and B5, the number of ‘extra’ species is considered as a proportion of the consumer species present at each time.

**Figure 4.**
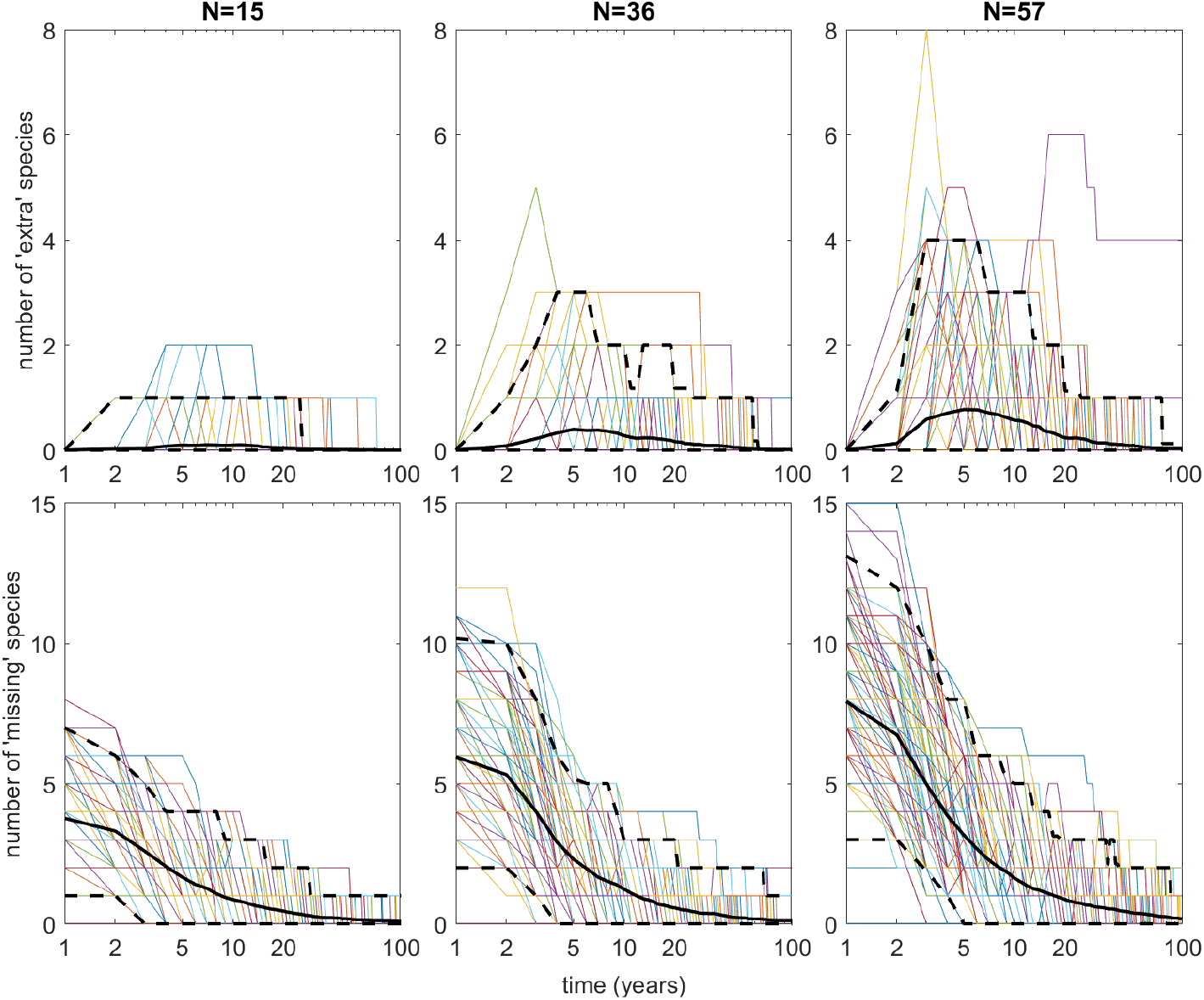
Number of ‘extra’ species and number of ‘missing’ species (relative to the endpoint community), over time. Time is plotted on a logarithmic scale. Each coloured line shows the results from one simulation; the black solid line shows the mean; the black dashed lines show the middle 95% of cases.

**Figure 5.**
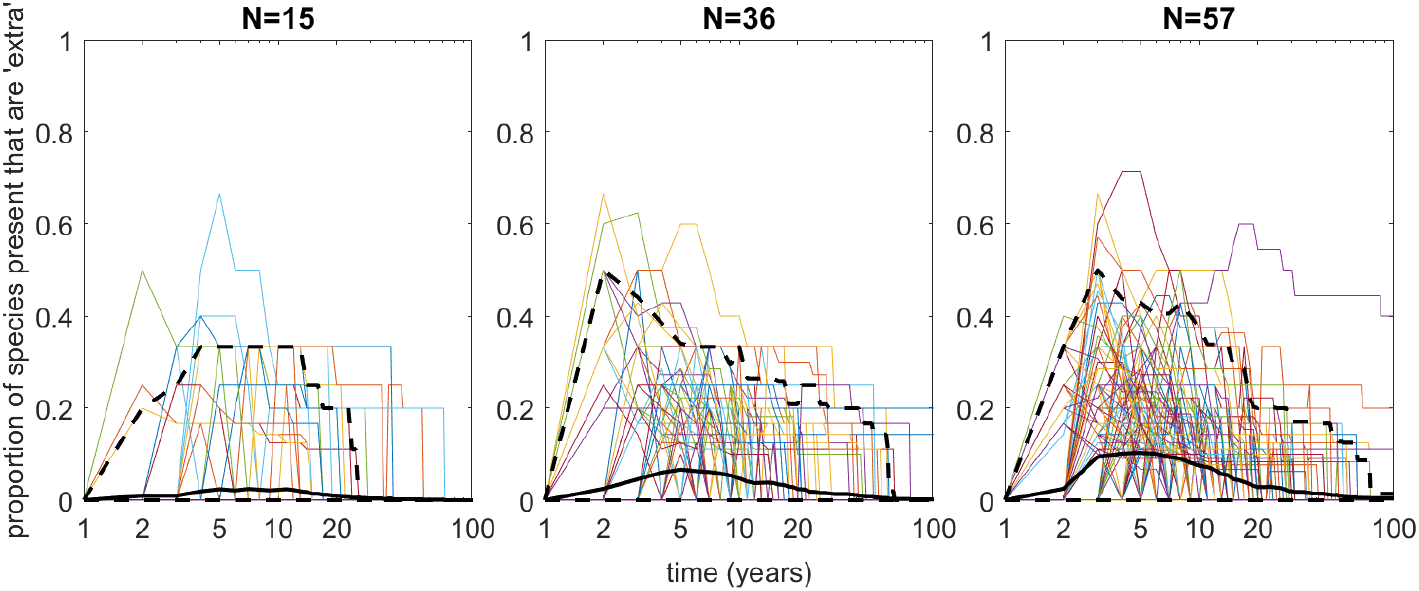
Proportion of the consumer species present that are ‘extra’, over time. Time is plotted on a logarithmic scale. Each coloured line shows the results from one simulation; the black solid line shows the mean; the black dashed lines show the middle 95% of cases.

In Figures 4 and B4, the black solid line shows the mean and the black dashed lines indicate the middle 95% of cases. The number of species ‘missing’ from the endpoint (i.e. permanent and uninvadable) community does, on average, decrease as the simulation goes on. The number of ‘extra’ species starts at zero, increases, and then decreases again, most often tending towards zero at long times. The results suggest that in most of our cases the number of species ‘missing’ and ‘extra’ tend to zero meaning that the simulated community does approach its predicted endpoint (i.e. permanent and uninvadable) set of species. However, in particular examples, the number of species ‘missing’ does not monotonically decrease but may go up as well as down (see Figures 4 and B4). Moreover there are cases where, even after 100 years, the simulation is not at its predicted endpoint community. Indeed, at time *t* = 100 years, over 13% of cases (211 out of 1577) still had at least one species either ‘missing’ or ‘extra’ relative to their endpoint community. These may be cases where the community dynamics are simply very slow, or may indicate cases where an alternative permanent and uninvadable set exists but has been missed by the method outlined in Section 2.2.2.

Figures 6 and B6 show how the number of cases with 0, 1, 2, 3 or 4 species ‘missing’ changes over time throughout the simulations, and likewise for the number of cases with 0, 1, 2, 3 or 4 ‘extra’ species. As an illustration of how these graphs are to be read, we see for example that for species pools of size 36, there were 172 cases that had at most one species ‘missing’ at *t* = 20 years, of which 109 cases had no species ‘missing’. Likewise, the graphs show that for the species pools of size 36, there were 188 cases that had at most one ‘extra’ species at *t* = 20 years, of which 166 had no ‘extra’ species. These values are out of a maximum of 192 cases (the species pools of size 36 that had a unique endpoint community).

**Figure 6.**
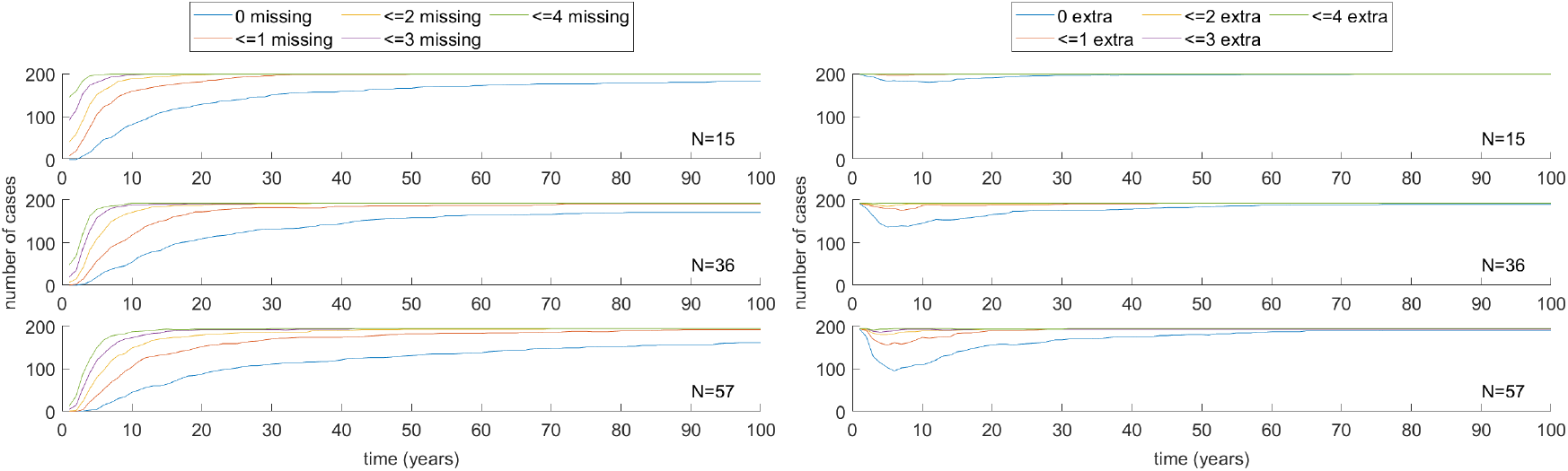
Left: Number of cases having no species ‘missing’ (blue), at most one species ‘missing’ (red), at most two species ‘missing’ (yellow), at most three species ‘missing’ (purple), and at most four species ‘missing’ (green); Right: Number of cases having no ‘extra’ species (blue), at most one ‘extra’ species (red), at most two ‘extra’ species (yellow), at most three ‘extra’ species (purple), and at most four ‘extra’ species (green). From top to bottom, the data are for *N* = 15, 36 and 57.

It is clear that by monitoring the community over the first ten years, it is not possible to deduce which species the final community will consist of. Indeed, 67% of our models (1061 out of 1577) still had species ‘missing’ after simulating for ten years, and over 22% of cases (355 out of 1577) had ‘extra’ species present that would not form part of the endpoint community.

The final question we had asked for the cases that had a unique endpoint community was, out of all the consumer species observed at any year throughout the simulation, what proportion of them were present in the endpoint community. The results, split up according to total species pool size, are given in Table 6. On average, this proportion decreased with an increasing species pool size, from 97% to 86%. It may well be lower than this for species pools with more than 56 consumer species.

**Table 6:** Proportion of the consumer species present in the community at any year that were also present in the unique endpoint community (average value given for each species pool size).

| N | 8 | 15 | 22 | 29 | 36 | 43 | 50 | 57 |
| --- | --- | --- | --- | --- | --- | --- | --- | --- |
|  | 0.9733 | 0.9677 | 0.9376 | 0.9286 | 0.9155 | 0.8883 | 0.8644 | 0.8682 |

## 4 Discussion

### 4.1 General discussion of results

### 4.2 Cases without a unique endpoint state

As well as the species pools where the method of Section 2.2.2 identified a unique permanent and uninvadable subset, there were 23 out of 1600 (1.4% of cases) for which the method did not find a unique endpoint community. Of these, there were 14 species pools (see Table 4) with two permanent and uninvadable subsets. In such cases the existence of more than one such subset suggests that the same species pool may lead to either one of these endpoint communities, depending on the initial densities of the species. In a small number of other cases (3 cases out of 1600), one permanent and uninvadable subset was found but it was not reliably reached by the sequential invasion method, while in 6 cases out of 1600 no permanent and uninvadable subset could be found at all. It is possible that these species pools could truly have no subsets that were permanent and uninvadable — they might, as in [8], reach a final stage of cycling among different communities, each of which can be invaded by some other species — or that all such subsets fail the (sufficient but not necessary) condition for permanence. Alternatively it might be that certain of their permanent and uninvadable subsets either take more than 200 iterations to reach or cannot be reached at all by the method of sequential invasion.

### 4.3 Limitations of methods

We used an ODE-based model in this study; each species density was a function only of time and we did not attempt to take into account spatial variations. Our model was also subject to a number of other limitations and caveats which should be remembered. Firstly, it is known that the Lotka-Volterra model might not represent realistic inter-species interactions. More complicated functional responses have been designed, but these too are just approximations, and they all require significantly more parameters to be chosen. Since these parameters are unknown, using a more complicated model would not add much realism to this study. Furthermore, the permanence condition that we used is specific to Lotka-Volterra equations [27, 28, 7, 8, 9] and no such criterion has been discovered for models with any other form of the species interactions.

The permanence criterion itself, based on a linear programming problem [27, 28], is very useful and considered fairly reliable; but it is only a sufficient condition, not a necessary one. It is possible for a subset *s* to be permanent and yet not possess an average Lyapunov function of the form (9). This means that this test could fail to recognise certain permanent subsets, which is another limitation to be kept in mind. Furthermore, we found that for large species pools (over about 25 species) it became very computationally intensive and time-consuming to compute the properties required for all possible subsets. Therefore we instead used a sequential invasion method for *N* > 25 which, while faster, has an increased potential to miss permanent and uninvadable subsets. Now, in addition to the ‘sufficient but not necessary’ issue, it could be that certain permanent and uninvadable subsets would either take more than 200 iterations to reach or cannot be reached at all by the method of sequential invasion — perhaps they could be reached if two or more species were allowed to invade simultaneously, but not if they invade one at a time.

A final aspect of the model that is likely to be unrealistic is that there is no time-dependence in the coefficients. In reality, there will be both seasonal variation over the course of each year, and changes in environmental conditions over a 100-year period. According to Hyndes et al. [30], climate-driven changes over a 100-year timescale are likely to affect both the production rates of temperate seagrasses and also the ecological interactions in seagrass communities. They predict that increasing temperatures over the next century are “likely to alter the number and identity of herbivorous species in temperate systems and increase consumption rates and lower the standing biomass of temperate seagrasses” [30]. Thus, a constant growth rate, constant carrying capacity, constant consumption rates, and so on, are unrealistic. However, as already discussed regarding the alternative inter-species responses, the inclusion of such time-dependent effects in the model would only be based on speculated values, and it would again preclude the use of the Lotka-Volterra permanence condition.

An extra caveat is that our models have only covered the scenario of ‘successful’ seagrass restoration. A similar method could also be used to simulate cases with only ‘partial success’ — with the seagrass biomass taking only a fraction of the values used here — and then compared with our own results.

## A Appendix Average Lyapunov function

For some choice of positive exponents *h*_*i*_ > 0, consider the function

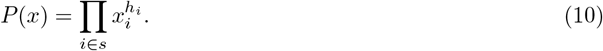

Note that *P* (*x*) = 0 on the boundary of the phase space for the subset *s*, and that *P* (*x*) increases as *x* moves away from the boundary into the interior. Note also that along an orbit *X*(*t*) satisfying (1),

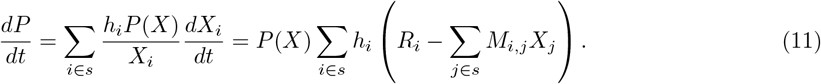

If the dynamics of the system meant that 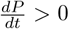 at all points near the boundary, then the boundary would be repelling orbits into the interior. In that case, *P* would be a Lyapunov function. However, as explained in [7, 8, 9], for the special Lotka-Volterra form of the equations (1), results of Hofbauer and Sigmund [27] and Jansen [28] mean that a weaker condition is sufficient for permanence. Namely, it is sufficient that *P* is an average Lyapunov function. For this we need only check that 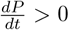 equilibria, which can be done by checking that

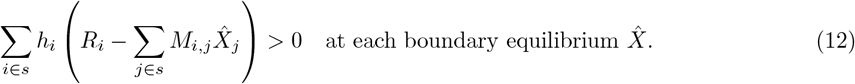

The quantities 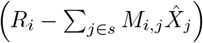 are proportional to the growth rates at the boundary equilibria. They are therefore zero for the species that have positive densities at the equilibria, but could have either sign for the absent species. The condition (12) therefore requires that the growth rates of the absent species have some weighted sum that is positive, and that the same weights (the *h*_*i*_) must work for all boundary equilibria 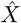.

Therefore, permanence can be established by solving the following linear programming problem [7, 8, 9]:

Choose *{z, h*_*i*_*}* to minimise *z* subject to:

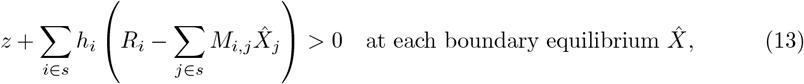

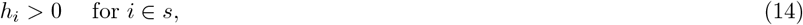

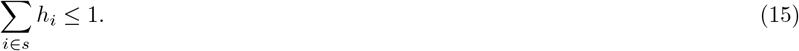

If *z <* 0 at this minimum, then by putting the corresponding values of the *h*_*i*_ into the formula (10) for *P* (*x*), we get an average Lyapunov function. Thus, if *z <* 0 at the minimum then the subset *s* does satisfy the permanence condition.

### B Supplementary Information

**Table B1:** Number of cases (out of 200) with at least one, and at least two, consumer species present.

| time period | N | 8 | 15 | 22 | 29 | 36 | 43 | 50 | 57 |
| --- | --- | --- | --- | --- | --- | --- | --- | --- | --- |
| $t \geq 2$ years | at least 1 consumer species | 52 | 67 | 67 | 90 | 73 | 75 | 90 | 89 |
|  | at least 2 consumer species | 11 | 20 | 31 | 56 | 41 | 61 | 63 | 70 |
| $t \geq 5$ years | at least 1 consumer species | 140 | 174 | 180 | 190 | 195 | 192 | 197 | 194 |
|  | at least 2 consumer species | 66 | 124 | 154 | 175 | 176 | 176 | 191 | 187 |
| $t \geq 10$ years | at least 1 consumer species | 183 | 197 | 200 | 199 | 200 | 200 | 200 | 200 |
|  | at least 2 consumer species | 122 | 174 | 182 | 193 | 195 | 197 | 199 | 199 |
|  | Out of total | 200 | 200 | 200 | 200 | 200 | 200 | 200 | 200 |

**Figure B1:**
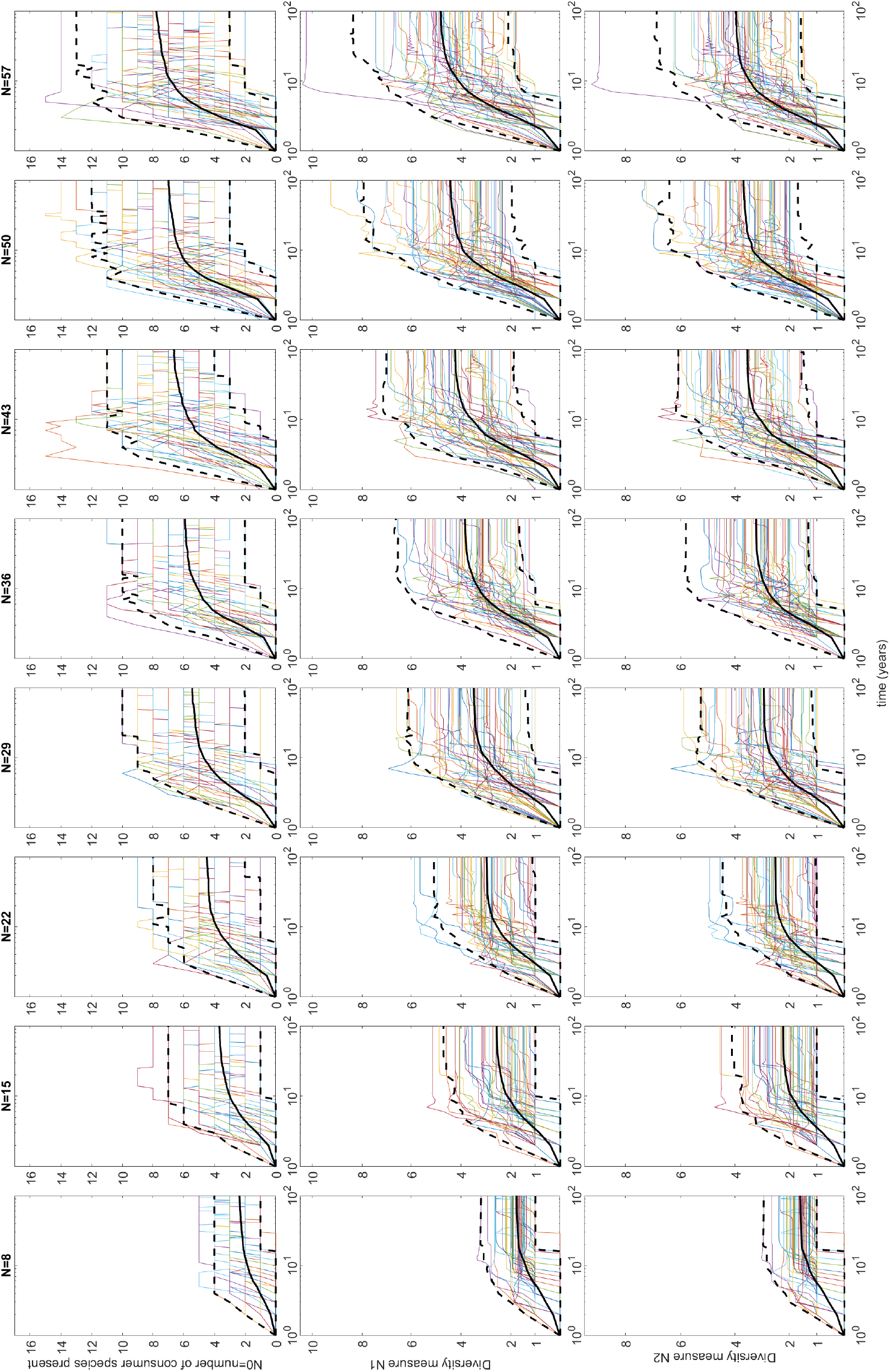
Diversity measures *N*_0_, *N*_1_ and *N*_2_ over time. Time is plotted on a logarithmic scale. For the purpose of plotting, we have set *N*_0_, *N*_1_ and *N*_2_ to be zero if no consumer species are yet present. Each coloured line shows the result from one simulation (for clarity only 50 out of each 200 are plotted). The black solid line shows the mean; the black dashed lines show the middle 95% of cases.

**Figure B2:**
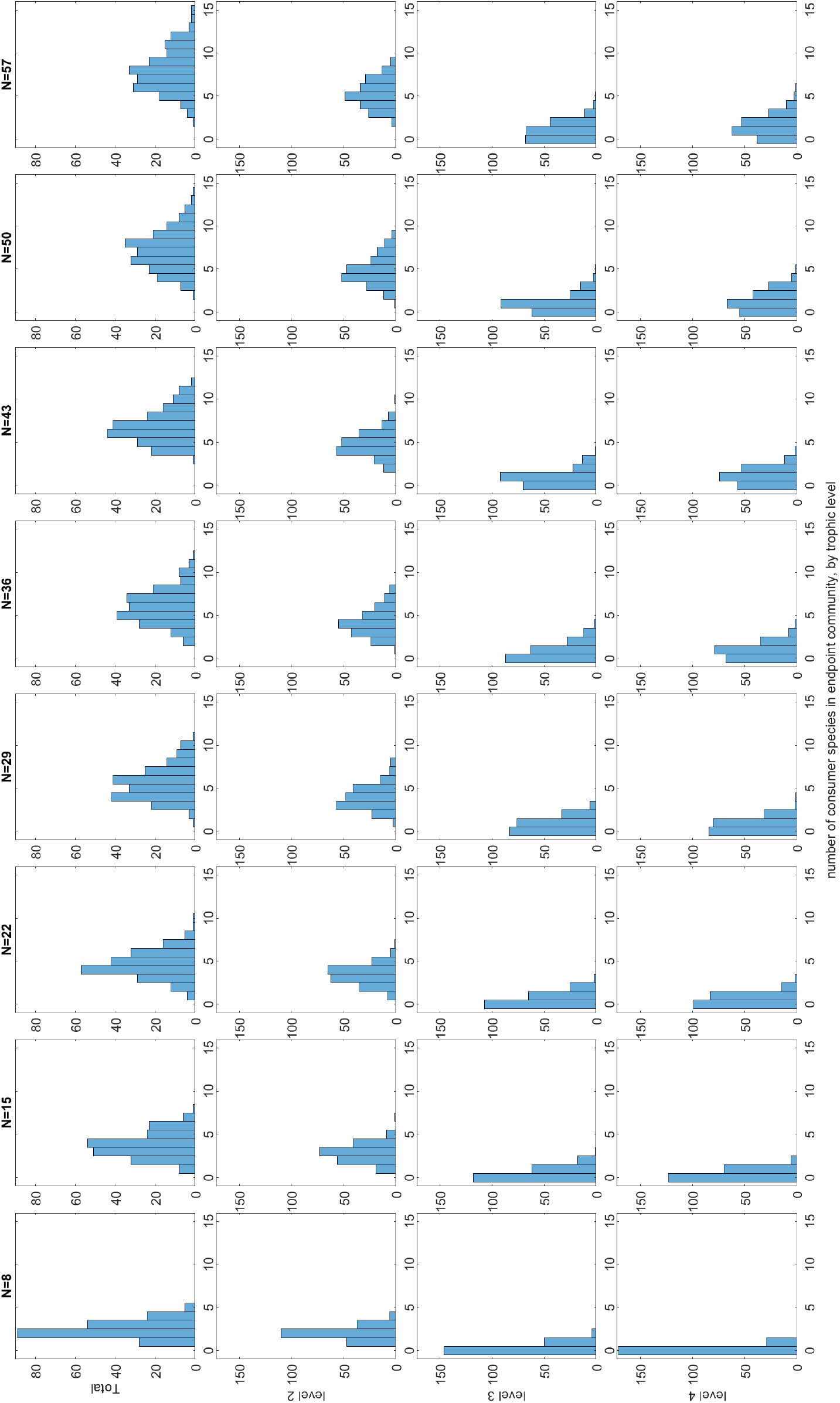
Number of consumer species in endpoint community, grouped by trophic level, for the different sized species pools.

**Figure B3:**
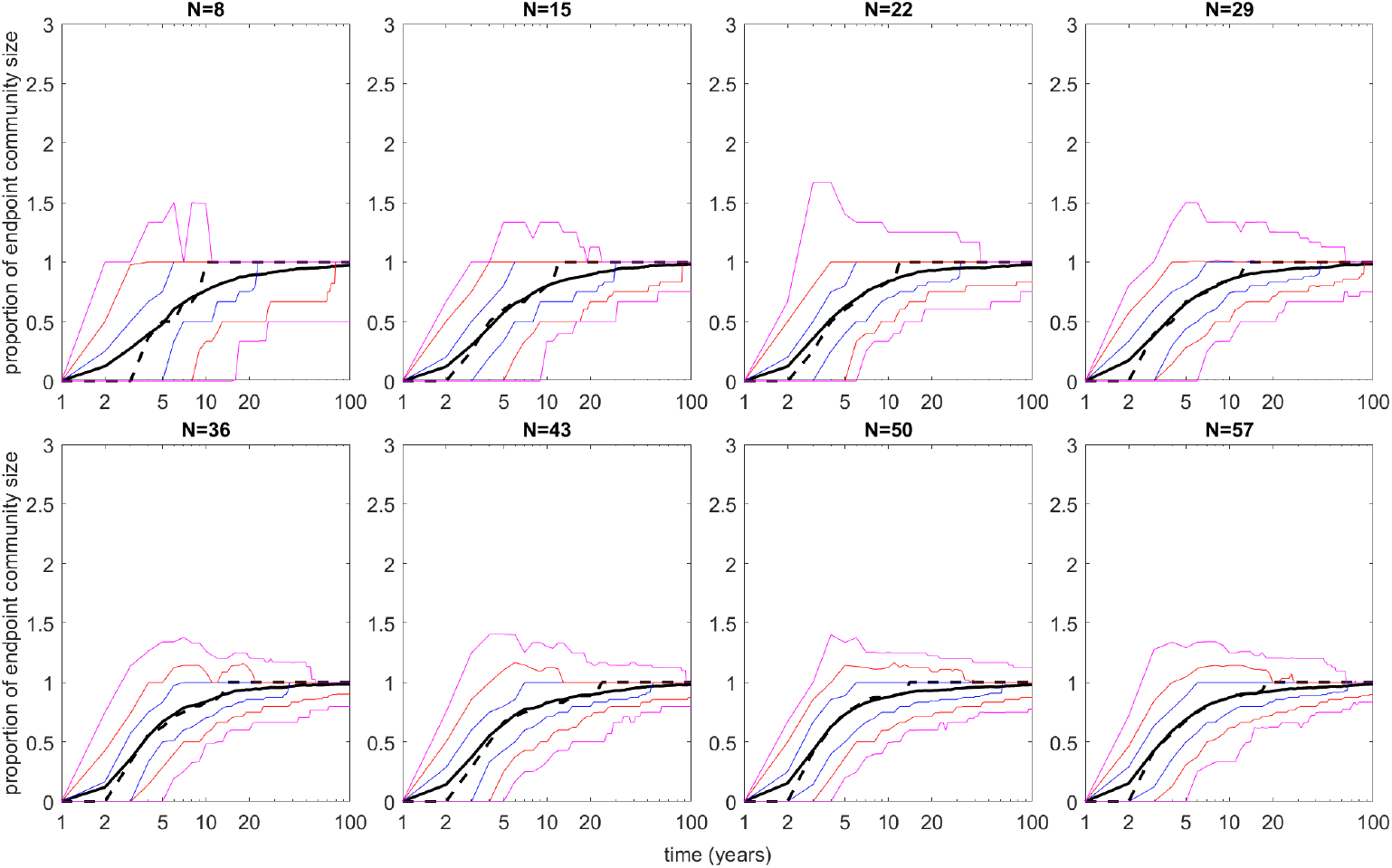
Number of consumer species present, as a proportion of the size of the endpoint community. The black solid line shows the mean and the black dashed line the median. The blue lines indicate the 25th and 75th percentiles, the red lines indicate the 10th and 90th percentiles, and the pink lines indicate the 2.5th and 97.5th percentiles. Time is plotted on a logarithmic scale.

**Figure B4:**
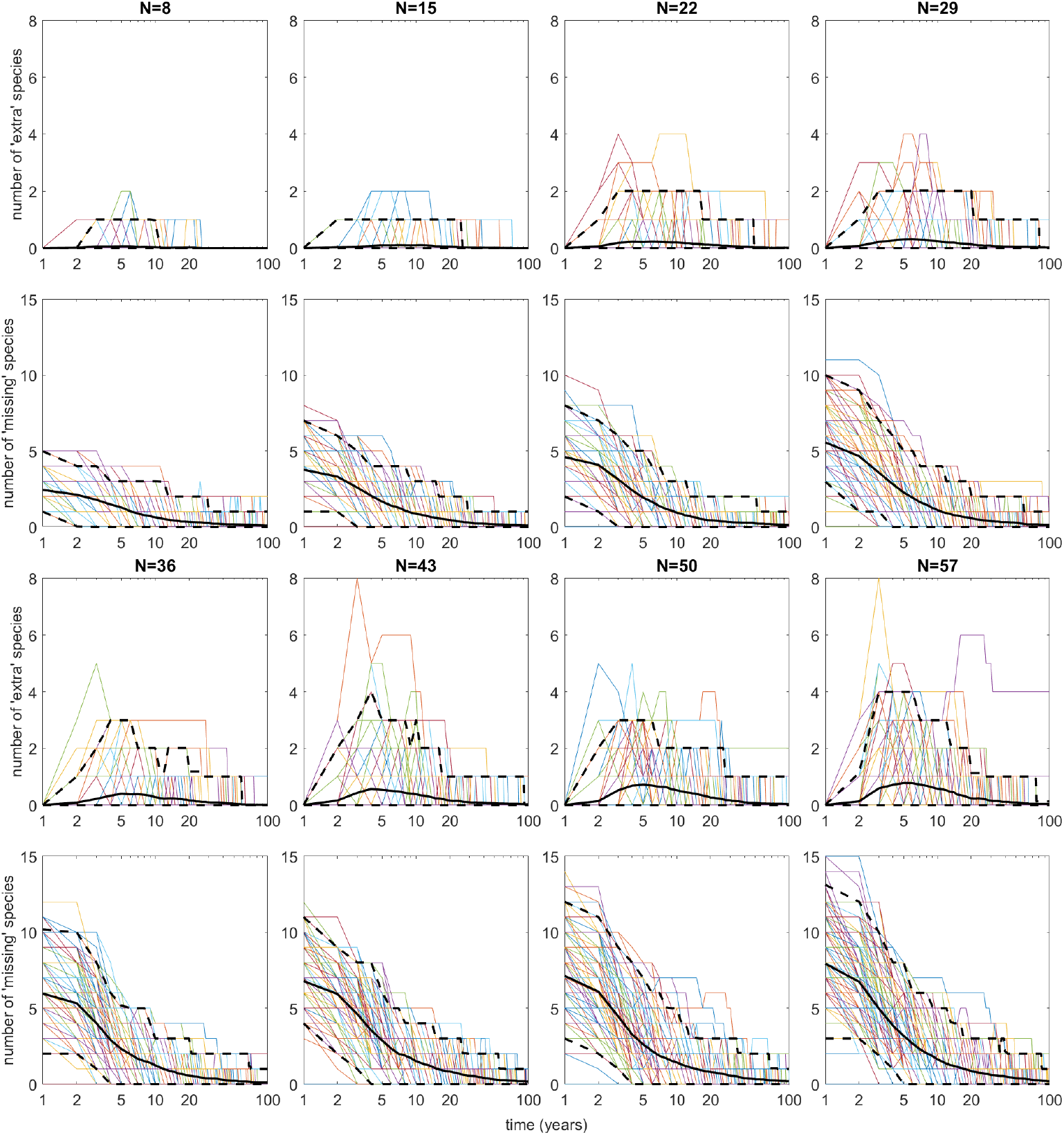
Number of ‘extra’ species and number of ‘missing’ species (relative to the endpoint community), over time. Time is plotted on a logarithmic scale. Each coloured line shows the results from one simulation; the black solid line shows the mean; the black dashed lines show the middle 95% of cases.

**Figure B5:**
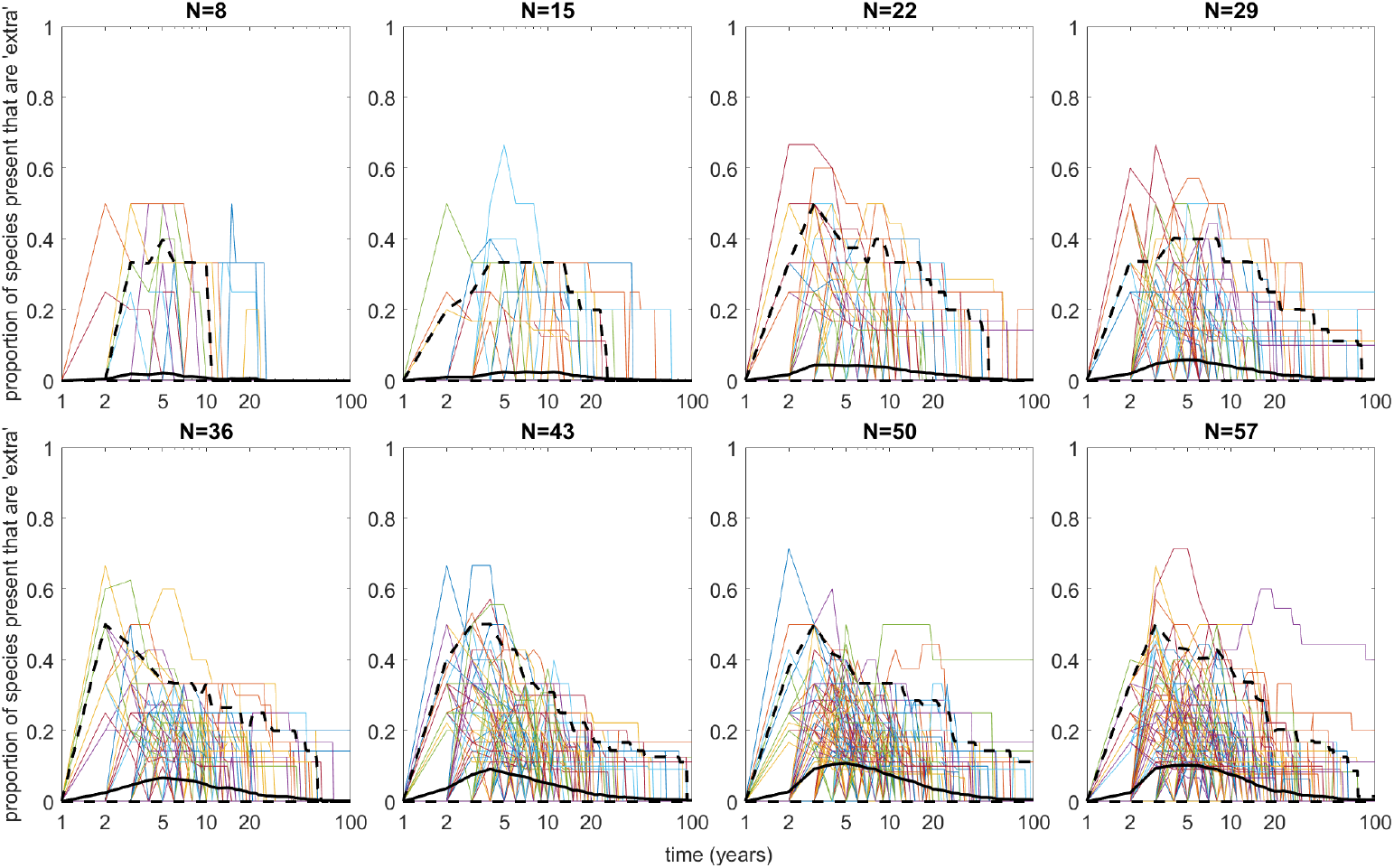
Proportion of the species present that are ‘extra’, over time. Time is plotted on a logarithmic scale. Each coloured line shows the results from one simulation; the black solid line shows the mean; the black dashed lines show the middle 95% of cases.

**Figure B6:**
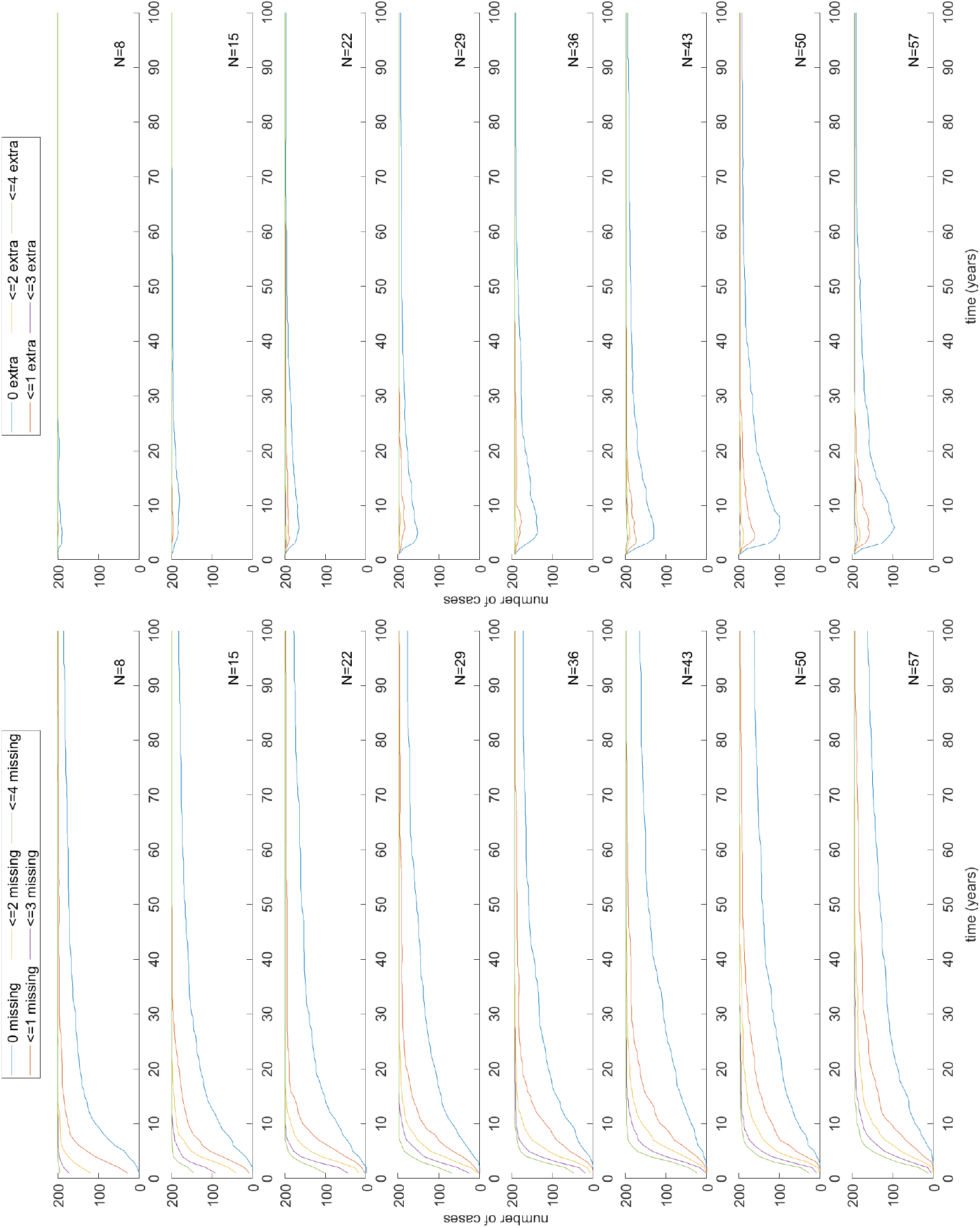
Left: Number of cases having no species ‘missing’ (blue), at most one species ‘missing’ (red), at most two species ‘missing’ (yellow), at most three species ‘missing’ (purple), and at most four species ‘missing’ (green); Right: Number of cases having no ‘extra’ species (blue), at most one ‘extra’ species (red), at most two ‘extra’ species (yellow), at most three ‘extra’ species (purple), and at most four ‘extra’ species (green). From top to bottom, the data are for *N* = 8, 15, 22, 29, 36, 43, 50 and 57.

## References

[1] Frederick T. Short, Blaine S. Kopp, Jeffery Gaeckle and Hitoshi Tamaki, Seagrass Ecology and Estuarine Mitigation: A Low-Cost Method for Eelgrass Restoration. Fisheries Science, 68(Sup2), pp 1759–1762 (2002).

[2] Robert J. Orth, Kenneth A. Moore, Scott R. Marion, David J. Wilcox and David B. Parrish, Seed addition facilitates eelgrass recovery in a coastal bay system. Marine Ecology Progress Series, Volume 448, pp 177–195 (2012).

[3] Marieke M. van Katwijk, Anitra Thorhaug, Núria Marbà, Robert J. Orth, Carlos M. Duarte, Gary A. Kendrick, Inge H. J. Althuizen, Elena Balestri, Guillaume Bernard, Marion L. Cambridge, Alexandra Cunha, Cynthia Durance, Wim Giesen, Qiuying Han, Shinya Hosokawa, Wawan Kiswara, Teruhisa Komatsu, Claudio Lardicci, Kun-Seop Lee, Alexandre Meinesz, Masahiro Nakaoka, Katherine R. O’Brien, Erik I. Paling, Chris Pickerell, Aryan M. A. Ransijn, Jennifer J. Verduin Global analysis of seagrass restoration: the importance of large-scale planting. Journal of Applied Ecology, 53, pp 567–578 (2016).

[4] Richard K. F. Unsworth, Chiara M. Bertelli, Leanne C. Cullen-Unsworth, Nicole Esteban, Benjamin L. Jones, Richard Lilley, Christopher Lowe, Hanna K. Nuuttila and Samuel C. Rees, Sowing the Seeds of Seagrass Recovery Using Hessian Bags. Frontiers in Ecology and Evolution, Vol. 7, 311, pp 1–7 (2019).

[5] C. Gamble, A. Debney, A. Glover, C. Bertelli, B. Green, I. Hendy, R. Lilley, H. Nuuttila, M. Potouroglou, F. Ragazzola, S. Rees, R. Unsworth and J. Preston, Seagrass Restoration Handbook. Zoological Society of London, UK (2021).

[6] Richard K. F. Unsworth, Leanne C. Cullen-Unsworth, Benjamin L. H. Jones, and Richard J. Lilley, The planetary role of seagrass conservation. Science, 377, pp 609–613 (2022).

[7] Richard Law and Jerry C. Blackford, Self-Assembling Food Webs: A Global Viewpoint of Coexistence of Species in Lotka-Volterra Communities. Ecology, Vol. 73, No. 2, pp 567–578 (1992).

[8] Richard Law and R. Daniel Morton, Alternative Permanent States of Ecological Communities. Ecology, Vol. 74, No. 5, pp 1347–1361 (1993).

[9] Richard Law and R. Daniel Morton, Permanence and the Assembly of Ecological Communities. Ecology, Vol. 77, No. 3, pp 762–775 (1996).

[10] Y. M. Tan, O. Dalby, G. A. Kendrick, J. Statton, E. A. Sinclair, M. W. Fraser, P. I. Macreadie, C. L. Gillies, R. A. Coleman, M. Waycott, K. van Dijk, A. Vergés, J. D. Ross, M. L. Campbell, F. E. Matheson, E. L. Jackson, A. D. Irving, L. L. Govers, R. M. Connolly, I. M. McLeod, M. A. Rasheed, H. Kirkman, M. R. Flindt, T. Lange, A. D. Miller and C. D. H. Sherman, Seagrass Restoration Is Possible: Insights and Lessons From Australia and New Zealand. Frontiers in Marine Science, 7:617 (2020).

[11] F. Kent, R. Lilley, R. Unsworth, S. Cunningham, T. Begg, P. Boulcott, C. Jeorrett, R. Horsburgh and M. Michelotti, Seagrass restoration in Scotland - handbook and guidance. NatureScot Research Report 1286 (2021).

[12] R. J. Orth, J. S. Lefcheck, K. S. McGlathery, L. Aoki, M. W. Luckenbach, K. A. Moore, M. P. J. Oreska, R. Snyder, D. J. Wilcox and B. Lusk, Restoration of seagrass habitat leads to rapid recovery of coastal ecosystem services. Science Advances, 6 (2020).

[13] Jurij Homziak, Mark S. Fonseca and W. Judson Kenworthy, Macrobenthic Community Structure in a Transplanted Eelgrass (Zostera marina) Meadow. Marine Ecology Progress Series, Vol. 9, No. 3, pp 211–221 (1982).

[14] Christoffer Boström and Erik Bonsdorff, Community structure and spatial variation of benthic invertebrates associated with Zostera marina (L.) beds in the northern Baltic Sea. Journal of Sea Research, 37, pp 153–166 (1997).

[15] A. L. Dale, R. McAllen and P. Whelan, Management considerations for subtidal Zostera marina beds in Ireland. Irish Wildlife Manuals, No. 28. National Parks and Wildlife Service, Department of Environment, Heritage and Local Government, Dublin, Ireland (2007).

[16] Allison L. Schmidt, Marta Coll, Tamara N. Romanuk and Heike K. Lotze, Ecosystem structure and services in eelgrass Zostera marina and rockweed Ascophyllum nodosum habitats. Marine Ecology Progress Series, Vol. 437, pp 51–68 (2011).

[17] Birgit Olesen and Kaj Sand-Jensen, Demography of Shallow Eelgrass (Zostera Marina) Populations — Shoot Dynamics and Biomass Development. Journal of Ecology, Vol. 82, No. 2, pp 379–390 (1994).

[18] Carlos M. Duarte and Carina L. Chiscano, Seagrass biomass and production: a reassessment. Aquatic Botany, 65, pp 159–174 (1999).

[19] Kaj Sand-Jensen, Biomass, net production and growth dynamics in an eelgrass (Zostera marina L.) population in Vellerup Vig, Denmark. Ophelia, 14:1–2, pp 185–201 (1975).

[20] Yi Zhou, Peng Liu, Bingjian Liu, Xujia Liu, Xiaomei Zhang, Feng Wang and Hongsheng Yang, Restoring Eelgrass (Zostera marina L.) Habitats Using a Simple and Effective Transplanting Technique. PLoS ONE, Vol. 9, Issue 4 (2014).

[21] Polly A. Penhale, Macrophyte-epiphyte biomass and productivity in an eelgrass (Zostera marina L.) community. Journal of Experimental Marine Biology and Ecology, Volume 26, Issue 2 pp 211–224 (1977).

[22] M. A. Borowitzka, P. S. Lavery and M. van Keulen, Epiphytes of seagrasses. In A. W. D. Larkum (Ed.), Seagrasses: Biology, Ecology and Conservation (pp441–462). Springer (2006).

[23] Owen L. Petchey, Anna Eklöf, Charlotte Borrvall and Bo Ebenman, Trophically Unique Species Are Vulnerable to Cascading Extinction. American Naturalist, 171, 5, pp 568–579 (2008).

[24] M. O. Hill, Diversity and Evenness: A Unifying Notation and Its Consequences. Ecology, Vol. 54, No. 2, pp 427–432 (1973).

[25] H. M. Anderson, V. Hutson and R. Law, On the Conditions for Permanence of Species in Ecological Communities. The American Naturalist, Vol. 139, No. 3, pp 663–668 (1992).

[26] R. Daniel Morton, Richard Law, Stuart L. Pimm and James A. Drake, On the models for assembling ecological communities. OIKOS, 75, pp 493–499 (1996).

[27] J. Hofbauer and K. Sigmund, The Theory of Evolution and Dynamical Systems: Mathematical Aspects of Selection. Cambridge University Press, Cambridge, England (1988).

[28] Wolfgang Jansen, A permanence theorem for replicator and Lotka-Volterra systems. Journal of Mathematical Biology, Vol. 25, pp 411–422 (1987).

[29] Nathaniel Virgo, Richard Law and Mark Emmerson, Sequentially assembled food webs and extremum principles in ecosystem ecology. Journal of Animal Ecology, 75, pp 377–386 (2006).

[30] Glenn A. Hyndes, Kenneth L. Heck, Jr., Adriana Vergés, Euan S. Harvey, Gary A. Kendrick, Paul S. Lavery, Kathryn McMahon, Robert J. Orth, Alan Pearce, Mathew Vanderklift, Thomas Wernberg, Scott Whiting and Shaun Wilson, Accelerating Tropicalization and the Transformation of Temperate Seagrass Meadows. Bioscience, 66, 11, pp 938–948 (2016).

